# Temporal profiling of the phosphate starvation response in Arabidopsis root hair cells reveals that induction of polycomb target genes does not depend on removal of H3K27me3 or H2A.Z

**DOI:** 10.1101/2024.07.14.603443

**Authors:** Dylan H. Holder, Roger B. Deal

## Abstract

Altered nutrient conditions can trigger massive transcriptional reprogramming in plants, leading to the activation and silencing of thousands of genes. To gain a deeper understanding of the phosphate starvation response and the relationships between transcriptional and epigenomic changes that occur during this reprogramming, we conducted a time-resolved analysis of transcriptome and chromatin alterations in root hair cells of Arabidopsis thaliana during phosphate (P) starvation and subsequent resupply. We found that 96 hours of P starvation causes induction or repression of thousands of transcripts, and most of these recover to pre-starvation levels within 4 hours of P resupply. Among the phosphate starvation-induced genes are many polycomb targets with high levels of H3K27me3 and histone variant H2A.Z. When induced, these genes often show increased H3K4me3 consistent with active transcription, but surprisingly minimal loss of H3K27me3 or H2A.Z. These results indicate that the removal of silencing marks is not a prerequisite for activation of these genes. Our data provide a cell type- and time-resolved resource for studying the dynamics of a systemic nutrient stress and recovery and suggest that our current understanding of the mechanisms for switching between silent and active transcriptional states is incomplete.

## Introduction

The nucleosome is the fundamental unit of chromatin in eukaryotes and is therefore a key substrate for transcriptional regulation. Nucleosomes can be modified via posttranslational modifications such as phosphorylation, ubiquitination, acetylation, methylation, or post-replicative exchange of canonical histone subunits with histone variants (Foroozani et al. 2022; Li et al. 2007). This array of modifications gives each nucleosome a unique identity thought to create a chromatin state that alters a given locus’s propensity for activation or silencing. It is known that proper organism development depends on the maintenance of these chromatin states across eukaryotes. Mutants deficient in several chromatin modification “writers”, “erasers”, and chromatin remodelers display pleiotropic developmental phenotypes that threaten the viability of the organism and its ability to properly respond to environmental cues (Talbert et al. 2019; Isbel et al. 2022). Terminally differentiated cells require dramatic changes in transcription rate at genes in response to intrinsic and extrinsic signals without cell division. Transcription machinery must therefore overcome the established chromatin state and transform the locus to accommodate gene activation or silencing. How the transcription machinery is able to rapidly overcome the repressive chromatin landscape at quiescent genes in response to activating signals, and then effectively re-silence these genes once those signals are gone, is not well understood.

Deposition of histone variant H2A.Z into postreplicated nucleosomes is required for proper development in eukaryotes. In animals, H2A.Z is required for embryonic viability and cancer cell suppression (Monteiro et al. 2014; Ávila-López et al. 2021; Clarkson et al. 1999; Faast et al. 2001). In plants, H2A.Z deficiency leads to severe post embryonic developmental phenotypes and impaired responses to abiotic stimuli (Noh and Amasino 2003; Deal et al. 2007; Sura et al. 2017; Smith et al. 2010; Kumar et al. 2010). Unlike H3K27me3 and H3K4me3, whose roles are characterized as repressors and activators respectively, H2A.Z enrichment at plant genes is found to either promote or repress transcription depending on the context (Cheng et al. 2019; Zhang et al. 2007; Foroozani et al. 2022). As a result, a unifying mechanism for H2A.Z’s role in regulating transcription has yet to be identified. H2A.Z is deposited by the SWR1 chromatin remodeling complex primarily into euchromatic genes and regulatory elements (Mizuguchi et al. 2004; Kobor et al. 2004). Genes occupied by H2A.Z nucleosomes in Arabidopsis are broadly defined by either a prominent enrichment peak near the transcription start site (TSS) or enrichment spanning the entire gene body. While TSS-proximal H2A.Z enrichment has been shown to promote both gene activation or repression depending on the experimental context, gene body enrichment is negatively associated with transcription, especially among genes implicated in environmental responses (Coleman-Derr and Zilberman 2012; Bönisch and Hake 2012; Weber et al. 2014).

Since the initial observation that gene body H2A.Z enrichment associates with transcriptionally repressed responsive genes, H2A.Z focused studies of osmotic, heat, light, phosphate, and hypoxic stress across *Arabidopsis thaliana, Oryza sativa, and Brachypodium distachyon* have implicated gene body H2A.Z as a repressor of environmentally responsive genes (Boden et al. 2013; Mao et al. 2021; Cortijo et al. 2017; Nguyen and Cheong 2018; Sura et al. 2017; Lee and Bailey-Serres 2019). While each study was conducted with varying techniques, tissues, and exposure lengths, two observations are consistent regardless of the stressor: 1) H2A.Z enrichment at these stress response genes is reduced following their induction. 2) These genes are often ectopically induced in H2A.Z-deficient plants, implicating H2A.Z as having a causal role in their repression. Gene body H2A.Z loss during activation, and overexpression in H2A.Z deficient plants, certainly implicates H2A.Z in having a causal role in gene repression. However, no unifying mechanism has been proposed to explain how H2A.Z mediated repression is achieved across all stress responses or how its eviction directly promotes transcription. Notably, the differences in primary structure between H2A.Z and canonical H2A have not been sufficient to explain the variant’s complex relationship with gene expression (Bönisch and Hake 2012).

A complete grasp of H2A.Z’s role in gene repression is likely not possible without understanding the surrounding chromatin environment into which it is deposited and observing changes during transcriptional activation or repression. One key factor to deciphering H2A.Z’s role in silencing is its relationship to the Polycomb Repressive Complex 1 and 2 (PRC1 and 2). The polycomb silencing system promotes gene silencing in both plants and animals and is hallmarked by trimethylation of histone H3 on lysine 27 (H3K27me3) by PRC2 and monoubiquitination of C-terminal lysine on H2A or H2A.Z (H2Aub) by PRC1 (Förderer et al. 2016). In mammalian stem cells, H2A.Z is associated with H3K27me3 at repressed genes (Ku et al. 2012; Wang et al. 2018). In *Arabidopsis,* H2A.Z and H3K27me3 enrichment are correlated across gene bodies and anticorrelated with transcript abundance. Furthermore, H2A.Z deficient plants have reduced H3K27me3 genome wide and PRC2 mutants have reduced H2A.Z, suggesting a mutual reinforcement of these two factors at target genes (Carter et al. 2018). However, recent work has called into question the silencing role of H3K27me3 at these genes and highlighted the requirement for H2A.Z monoubiquitination by PRC1 in plant polycomb gene silencing (Gómez-Zambrano et al. 2019).

During a phosphate starvation response (PSR), plants undergo system wide transcriptional reprogramming to improve inorganic phosphate uptake efficiency from the soil and internal P utilization. This involves induction and repression of transcripts from a diverse set of biological processes including phosphate transport (intercellular, extracellular, and intracellular), phosphatase secretion in root, supplementing phospholipid loss in the plasma membrane, ATP conservation, and prominent reorganization of root system morphology (Yang et al. 2024). The phosphate starvation response has been characterized at the transcriptome level in whole shoot and root (Thibaud et al. 2010; Shukla et al. 2021; Misson et al. 2005). However, how a phosphate starvation-induced (PSi) gene’s chromatin state changes to allow for this responsive activation and later re-silencing is not understood. Root hair cells are among the first cells to encounter changes in phosphate availability in the soil, as they are located on the root epidermis and project into the substrate in search of nutrients. As a terminally differentiated cell type, they possess their own unique chromatin landscape that is a history of the cell’s journey through differentiation and dictates transcription events to come as the cell encounters exogenous challenges.

Numerous PSi genes are marked with H3K27me3 along with H2A.Z under normal conditions, and H2A.Z deficient plants overexpress several key PSi genes even on P-rich media (Smith et al. 2010; Coleman-Derr and Zilberman 2012; Carter et al. 2018). Phosphate (P) is a growth limiting nutrient and plants must rapidly overcome these repressive chromatin marks to activate and then resilience PSi genes to adapt to changing P availability in the soil. Given H2A.Z’s relationship with polycomb mediated silencing and its implication in responsive gene repression, we hypothesized that H2A.Z-containing nucleosomes mediate the endogenous rate of gene activation and aid in rapid re-silencing through cooperation with polycomb, possibly acting as a preferred substrate for polycomb machinery promoting a more flexible alternative to repression via DNA methylation. To test this hypothesis and explore the dynamics of PSi gene induction and subsequent resilencing, we performed a cell-type and time resolved multi-omic study of transcript levels, histone modifications, and chromatin accessibility in Arabidopsis. We probed the chromatin and transcriptional landscape over a P starvation and resupply time course experiment to understand how root hairs overcome the repressive effects of H2A.Z and H3K27me3. We took rapid measurements in root hair cells following P resupply to examine the order of events that take place on chromatin to establish an H2A.Z-mediated silent state. To address the confounding factors inherent with traditional whole tissue chromatin profiling techniques, we utilized Isolation of Nuclei TAgged in a specific Cell Types (INTACT) followed by Cleavage Under Targets and Tagmentation (CUT&Tag) to capture the chromatin landscape exclusively in the root epidermal hair cell, where P sensing and uptake first occurs (Kaya-Okur et al. 2019; Deal and Henikoff 2010).

To our surprise, despite robust PSi gene induction and re-silencing across the time course, H2A.Z and H3K27me3 dynamics were modest throughout the time-course, with both being maintained at high levels across gene bodies after gene induction. We went on to find that while H3K27me3 is highly enriched at PSi genes, it does not appear to be required for P dependent gene activation or re-silencing. The data presented here provide a resource for studying *in-vivo* cell type-specific transcriptome and chromatin dynamics during a systemic transcriptional activation and re-silencing event. The results not only provide a resource for more deeply understanding the PSR, but also indicate that transcription can occur even in the continued presence of H3K27me3 and H2A.Z across gene bodies.

## Results

### Transcript profiling during phosphate starvation and resupply reveals major starvation-induced changes that generally recover rapidly upon resupply

To examine transcriptional changes associated with P starvation and resupply, we utilized an Arabidopsis INTACT transgenic line in which nuclei of the root hair cell type were labeled (Deal and Henikoff 2010). We chose the root hair cell type due to their physical interaction with the phosphate-containing soil environment and their pronounced morphological changes in response to P starvation. Measuring this response in a single cell type also eliminates the confounding factors inherent to probing the transcriptional landscape across the diverse population of cell types that compose the root. Seedlings were grown vertically on ½ MS plates for 10 days (0h+P), transferred to phosphate depleted media for 96 hours (96h-P), and finally moved back to P rich media and sampled at 4 time points following P resupply (30m, 1h, 4h+P) (**Figure 1a**). We settled on this sampling scheme given that initial RT-qPCR experiments on whole roots with higher temporal resolution showed that two hallmark PSR genes are induced after 96h -P and reach near-complete repression within 4 hours of P resupply (**Figure 1b**). At each of the 6 time points sampled, root segments containing fully differentiated root hair cells (∼1 cm segment from above the differentiation zone and below the first lateral roots) were harvested for INTACT nuclei purification (Wang and Deal 2015) followed by nuclear RNA-seq and ATAC-seq analyses.

**Figure 1.**
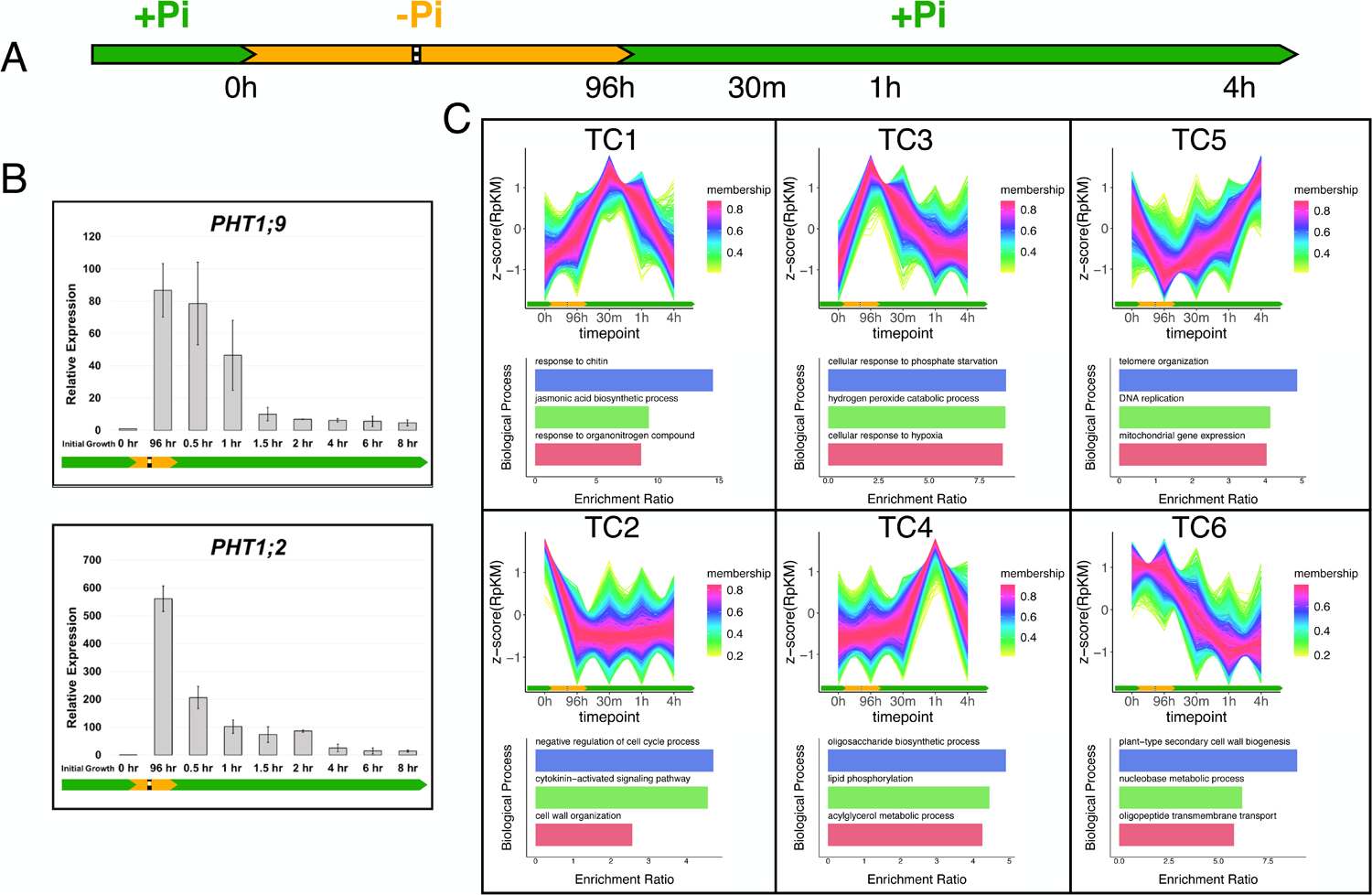
Root-hair RNA-seq during a phosphate starvation reveals several classes of phosphate starvation responses. (**A**) Schematic of the phosphate (P) starvation and resupply treatment. Each time point represents a sample taken for INTACT-RNA-seq. (**B**) Quantitative RT-PCR analysis of RNA from whole roots exposed to P starvation and resupply. Error bars represent standard deviation from the mean for 3 replicates. Expression is represented as amounts relative to the pre-starvation time point (0h). (**C**) C-means clustering analysis of INTACT-RNA-seq results via TC-seq. Each line represents a single differentially expressed gene. Line color corresponds to the degree of membership to each c-means cluster. Below each cluster are enrichment ratios for three biological processes enriched in each cluster as identified by GO analysis via WebGestalt.

Differentially expressed genes (DEGs) were c-means clustered into 6 groups based on common expression patterns across the time-course via TC-seq (**Fig 1c, Dataset S1**) (Mengjun 2019). This analysis revealed 5,674 genes differentially expressed between any two measured timepoints, with 2,592 differentially expressed genes after 96h of P starvation (**Dataset S1**). We deemed the 915 genes in time course cluster 3 (TC3) as phosphate starvation-induced (PSi) genes due to increased transcript after 96 hr P starvation that falls rapidly after P resupply (TC3, **Figure 1c**). Gene ontology analysis confirmed that known PSi genes occupy this cluster (including *PHT1;2* and *PHT1;9*) with enrichment for protein families like secretory peroxidases and hypoxia response genes which have been identified in other PSR experiments (**Dataset S2**). The P1BS motif, known to be bound by the PSR transcription factor (TF) PHR1 and related TFs, was enriched in the accessible chromatin (via INTACT-ATAC-seq) overlapping with TC3 promoters when compared to the other TC clusters (**Dataset S3**, p-value = 4.15e-5, enrichment ratio 2.65). Additionally, Simple Enrichment Analysis of ATAC-accessible sites in TC3 promoters for known TF motifs revealed enrichment for several MYB family TF motifs involved in biological processes known to be linked with phosphate sensing including cell polarity, nitrate sensing, circadian clock, and PHR1-like transcription factors (**Dataset S3**).

Interestingly, we also identified several other classes of transcriptional responses during the starvation/resupply event (**Figure 1c, Figure S1**). Of note, TC5 reflects a group of phosphate starvation-repressed (PSr) genes that recover to baseline levels after 4 h of P resupply. These genes are enriched for functions involved in maintaining chromosome integrity and mitochondrial gene expression, possibly reflecting transcriptional reprioritization as ATP production falls during starvation. TC2 genes are also silenced during P starvation but fail to recover within 4 h of resupply. These consist of cell cycle and cell wall organization genes, likely reflecting the cellular expansion root hair cells undergo during P starved conditions. A unique pattern of expression is observed in the TC4 genes, as these are maximally induced within 1 hour of P resupply but then return to baseline by 4 hours. These data are a rich resource to explore the biological processes involved in the PSR that do not follow the canonical path of upregulation during starvation and repression during resupply.

### INTACT-CUT&Tag recapitulates ChIP-seq profiles

To examine chromatin dynamics across the time course, we next used INTACT-purified root hair cell nuclei for CUT&Tag on H3K4me3, H3K27me3, and H2A.Z. K-means clustering of the root-hair CUT&Tag datasets with corresponding publicly available ChIP-seq from seedling tissue at genes for each mark revealed a strong agreement across the different data types (**Figure 2a**). Peak calling using MACS2 revealed a high degree of overlap between whole seedling and root hair nuclei across H3K27me3, H2A.Z and H3K4me3 (50, 75, and 70% of root hair peaks overlap with seedling peaks respectively) (**Figure 2b**) (Feng et al. 2012). However, root hair cells still possess a unique chromatin footprint when compared to seedling tissue with 10945, 3492, and 4807 root hair specific peaks for H3K4me3, H3K27me3, and H2A.Z respectively (**Figure 2c**). Peaks called exclusively in ChIP samples still have enrichment in the corresponding CUT&Tag sample, likely reflecting limitations of using a peak caller designed for high background ChIP-seq data on low background CUT&Tag data. Stratifying genic chromatin profiles by their transcript levels in the root hair RNA-seq data recapitulates the previously observed relationship between each chromatin mark, ATAC accessibility, and expression. For example, H3K27me3 is negatively correlated with transcript abundance, H3K4me3 is positively correlated with transcript abundance with enrichment downstream of the TSS, while H2A.Z enrichment shifts from the TSS to the gene body as transcript abundance falls (**Figure 2d**). The observable differences in enrichment across expression quantiles also confirm that this technique has a dynamic range comparable to other chromatin profiling techniques and will reliably measure changes in enrichment at a given site across a P starvation event.

**Figure 2.**
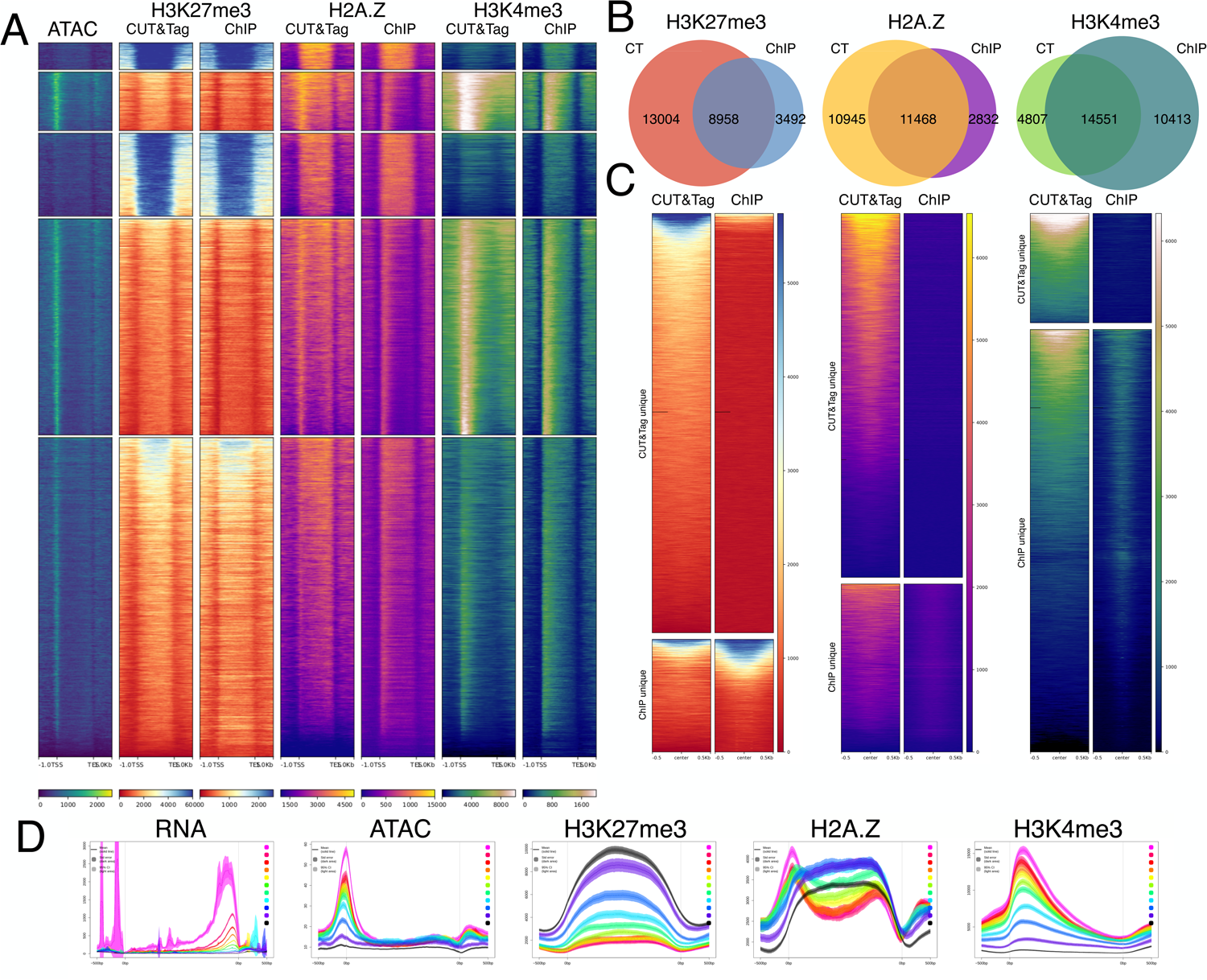
Root-hair INTACT-CUT&Tag compared to seedling ChIP-seq. (**A**) K-means clustered heatmaps of RPKM normalized signal at all Arabidopsis genes (TSS to TES, +/− 1 kb) comparing the enrichment of H3K27me3, H2A.Z, H3K4me3 between whole seedling ChIP-seq samples and root hair cell CUT&Tag samples. (**B**) Overlapping peaks between whole seedling ChIP-seq samples and root hair CUT&Tag samples for H3K27me3, H2A.Z, and H3K4me3. (**C**) RPKM normalized heatmaps of seedling ChIP-seq unique peaks or root hair CUT&Tag unique peaks (+/− 0.5 kb from center). **(D**) RPKM normalized average plots for each signal stratified by RNA expression decile in root hair cells showing highest expressed (warm colors) to lowest expressed (cool colors) genes. Black represents non-expressed genes.

#### Chromatin dynamics across the time course are mostly limited to H3K4me3 changes

Although each chromatin profile correlated with steady state transcript levels as expected, we found this did not hold when comparing the dynamics of these profiles across time. After 96h -P, only 51 genes were differentially enriched for H3K27me3 and only 9 genes were differentially enriched for H2A.Z (padj<.05) (**Figure 3a**). The lack of H2A.Z and H3K27me3 dynamics is also observed across all differentially expressed genes, where log2 fold changes are restricted to between +/− 1 (**Figure 3b**). H3K4me3 changed the most across the P starvation and repletion, with 1,900 genes differentially enriched for the mark after 96h -P. However, only 390 of those genes overlapped with differentially expressed genes across the same time points (**Figure 3a, Figure S2**). While accessibility changes are observed at DEGs across the time course, almost no changes were statistically significant (**Figure 3b, Figure S2**). This lack of significant ATAC changes is true both at DEG promoter regions (−200bp +100bp TSS) and at ATAC hypersensitive sites and is in contrast to a previous study that found thousands of differentially accessible regions after 10 days of phosphate starvation in whole roots (Barragán-Rosillo et al. 2021). The discrepancy between this study and our findings could be explained by the experimental conditions. While Barragán-Rosillo et al. collected whole roots from 10 day old seedlings grown from germination on phosphate limited media, we collected root hair cell nuclei grown for only 96h on phosphate depleted media. Our results suggest that significant changes in chromatin accessibility may depend on longer starvation periods or even cell division. We next examined the persistence of each chromatin mark across the time course to determine the stability of the chromatin changes caused by 96h of P starvation (**Figure 3a**). For H3K4me3, 79% of the 1900 genes with differential H3K4me3 enrichment at 96h were still differentially enriched after 4h of P resupply compared to 0h. Likewise for H3K27me3, 63% of the 51 differentially enriched genes after starvation are still differentially enriched after 4h of resupply. While only 9 genes were differentially enriched for H2A.Z, 3 of those were still differentially enriched after 4h. Interestingly, the RNA landscape returns to pre-starvation levels faster than the measured chromatin marks, with 74% of the 2592 differentially expressed transcripts after starvation returning to pre-starvation levels after 4h of resupply.

**Figure 3.**
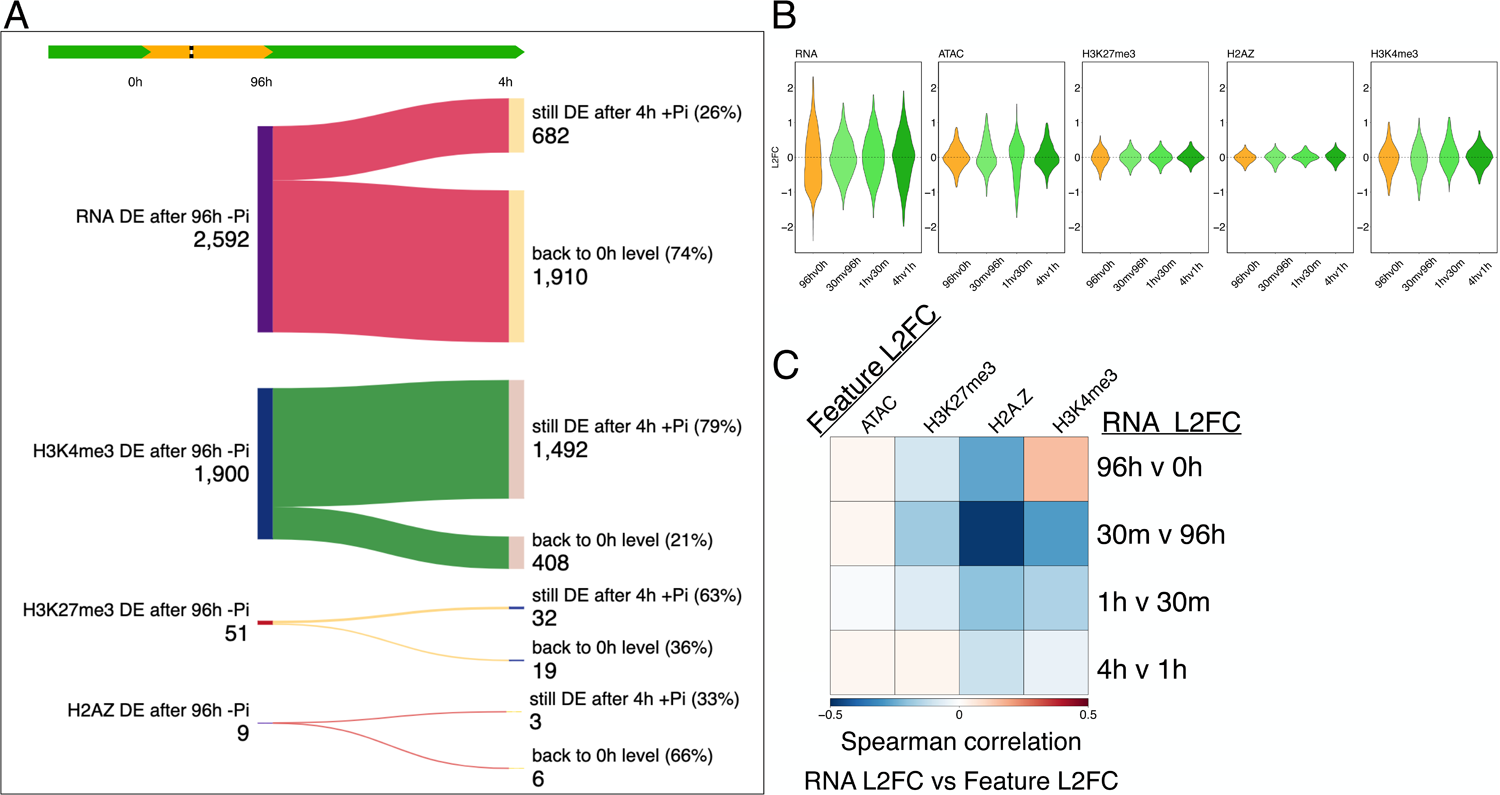
Chromatin dynamics during a phosphate starvation and recovery. (**A**) Flow chart of each chromatin profile tracking the trajectory of each differentially enriched (DE) gene at 96h -P to 4h of P resupply (padj < 0.05 via DESeq2). Generated using SankeyMATIC. (**B**) Log2 fold change (L2FC) violin plots of each chromatin profile across the time course. Data in the plot are limited to differentially expressed genes identified via TC-seq (n= 5,647) with outliers excluded from each plot. (**C**) L2FC Spearman correlation between RNA and each chromatin profile across the time course. Each L2FC is measured relative to the previous time-point.

To better understand how each chromatin profile changes in relation to transcript abundance we performed Spearman correlations between the Log2 fold-change (L2FC) for each mark at each gene (relative to the previous time point) and the corresponding L2FC of RNA at each gene (**Figure 3c**). We found no correlation between RNA changes and chromatin accessibility changes measured by ATAC, and weak negative correlations between changes in RNA and both H2A.Z and H3K27me3. That said, changes in H2A.Z at genes are still anticorrelated with RNA dynamics across the time course, perhaps indicating that H2A.Z dynamics reflect general transcription as opposed to influencing a specific mechanism of transcription at certain genes (**Figure 3c**). Interestingly, H3K4me3 dynamics were positively correlated with RNA level changes after 96h of P starvation, but not between any of the shorter time points after P resupply (**Figure 3c**).

### Many phosphate starvation-induced genes are enriched for gene body H2A.Z and H3K27me3 but these marks are not significantly altered after 96h of P starvation

Given the presumed role of H2A.Z in repression of responsive genes, we sought to identify a chromatin signature that might distinguish the responsive gene clusters identified by TCseq (**Figure 1c**). In particular, we expected many PSi genes to be in a chromatin state with enrichment for H3K27me3 and H2A.Z in the gene body (gbH2A.Z) before starvation (0h). Upon visualization using deepTools, we confirmed that the PSi genes in TC3 are in fact enriched for H3K27me3 compared to other clusters and contain gbH2A.Z (**Figure 4a**). Considering the high average of H3K27me3 and H2A.Z gene body enrichment at TC3, it was surprising to discover that genes in TC3 also have the highest average transcript abundance before P starvation, along with TC6 (**Figure 4a**).

**Figure 4.**
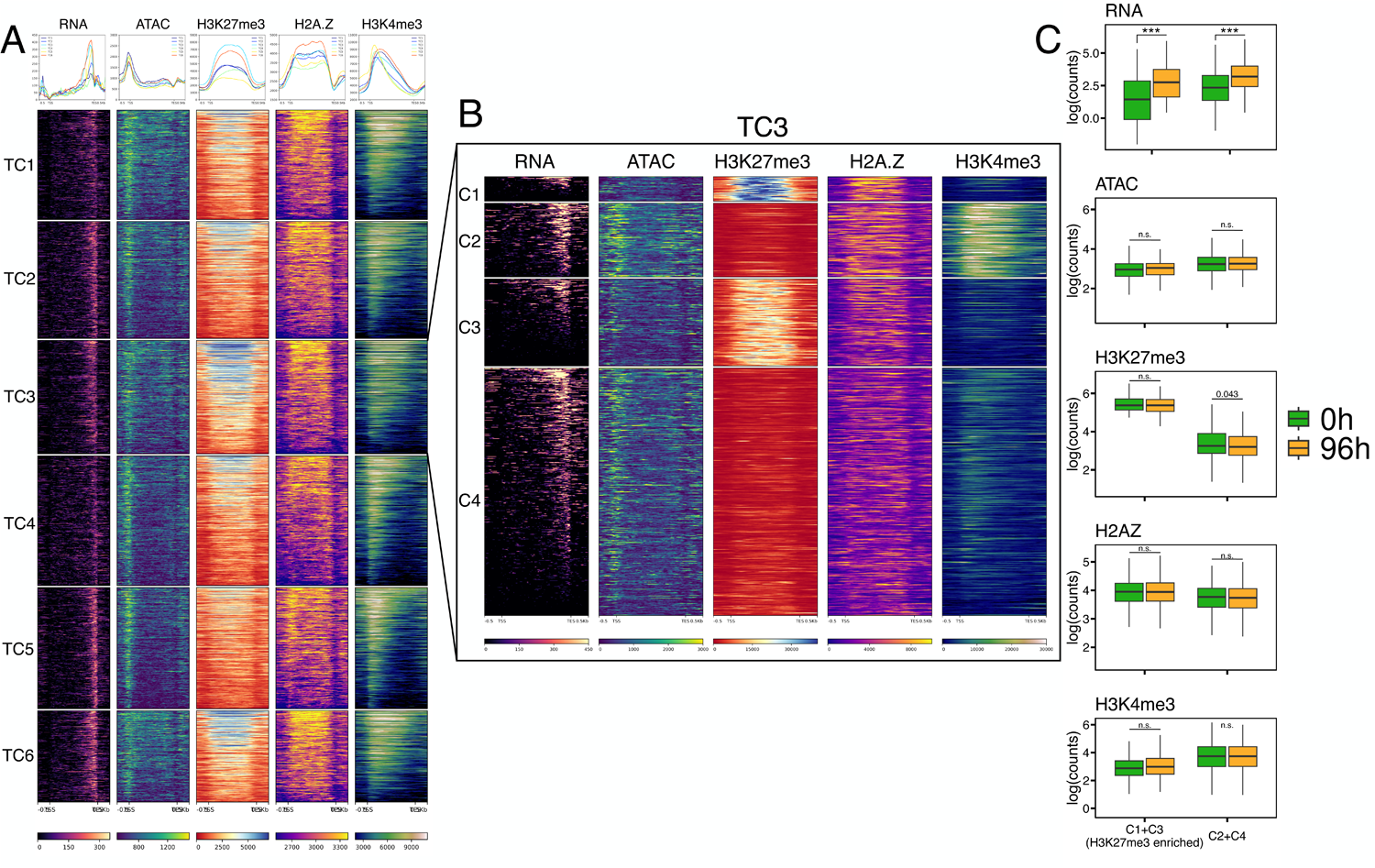
Phosphate starvation-induced (PSi) genes are enriched for H3K27me3 and gbH2A.Z but do not show significant changes in chromatin profile enrichment after P starvation. (**A**) RPKM normalized heatmaps of each RNA-seq TC cluster before starvation (0h), ranked by average signal across all profiles within each cluster. (**B**) K-means clustered heatmaps of chromatin profiles at all TC3 PSi genes (+/− 0.5 kb) before P starvation (0h) in root hair cells, RPKM normalized. Ranked by RNA signal. (**C**) Boxplot of log(quantile normalized counts) for each profile at H3K27me3 enriched TC3 genes (C1 and C3) and non-H3K27me3 enriched TC3 genes (C2 and C4) at 0h and 96h. ***p<0.001 (unpaired Student’s t-test).

Upon closer examination by k-means clustering, TC3 has more transcriptionally quiescent H3K27me3 and gbH2A.Z enriched genes but also has active genes with more RNA abundance than other clusters (**Figure 4b**). This indicates that while PSi genes have a wide spectrum of steady state transcript levels, many of these genes are indeed silent under normal conditions and are enriched for H3K27me3 and gbH2A.Z. We isolated H3K27me3 enriched genes (C1 and C3) from TC3 via k-means clustering and compared them to the remaining genes in TC3 (C2 and C4). We found that H3K27me3 enriched genes in TC3 had lower ATAC enrichment, H3K4me3 enrichment, and transcript abundance (**Figure 4b**).

We hypothesized that since H2A.Z at these genes is required for their repression under P rich conditions (Smith et al. 2010), we would observe an eviction of H2A.Z and H3K27me3 after 96 hr P starvation and a re-deposition of H2A.Z and H3K27me3 during their re-silencing, based on their established co-dependence (Carter et al. 2018; Smith et al. 2010). Surprisingly, PSi genes with the highest transcriptional dynamics displayed only minor changes in H3K27me3 and H2A.Z (**Figure 4c**). This increase in PSi transcript and maintenance of the H3K27me3/gbH2A.Z chromatin state during P starvation were confirmed by RT-qPCR and ChIP-qPCR on whole root tissue (**Figure S3**).

### Phosphate starvation-induced genes are highly enriched for H3K27me3, but PRC2 activity is not required for their phosphate-dependent activation or silencing

To confirm that the elevated enrichment of H3K27me3 was unique to PSi genes and not just a reflection of the expression levels of the genes at 0h, we generated a control dataset of non-differentially expressed genes with equivalent expression to each gene in TC3 at 0h -P. Comparing H3K27me3 enriched PSi genes to the control set reveals that they have significantly greater enrichment of H3K27me3 (**Figure 5a**). Interestingly, these H3K27me3 enriched PSi genes have more H3K27me3 on average than even the lowest 10% of expressed genes (**Figure 5b**).

**Figure 5.**
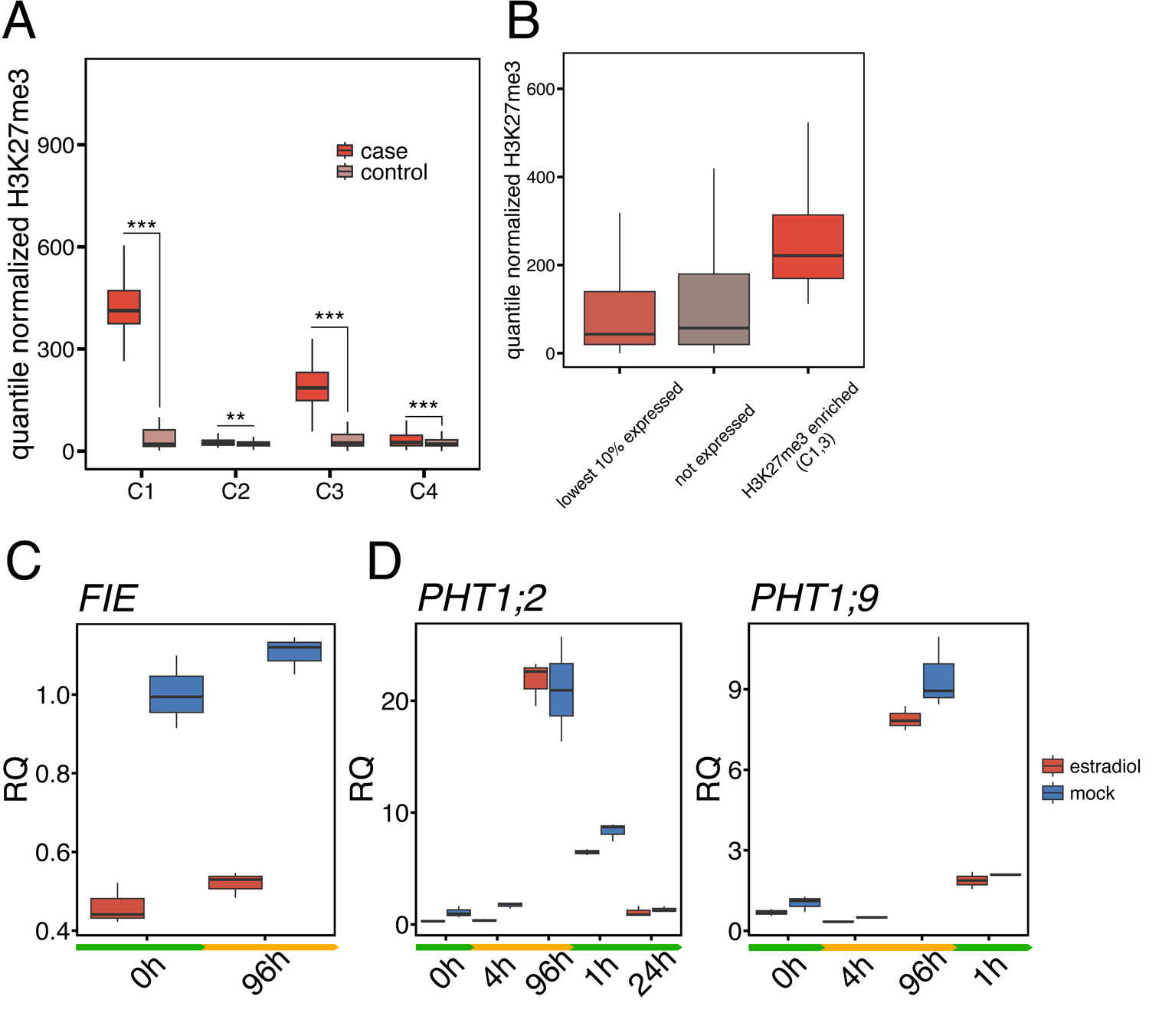
H3K27me3 enriched PSi genes do not rely on PRC2 for proper regulation. (**A**) Boxplot of quantile normalized H3K27me3 enrichment at each k-means cluster from A compared to a set of control genes with equivalent expression at 0h. ***p<0.001 (Wilcoxon two-sided test). (**B**) Boxplot of quantile normalized H3K27me3 enrichment at the lowest 10% of expressed genes, not expressed genes and H3K27me3 enriched genes from TC3 (C1 and C3 from Figure 4b). (**C**) Relative quantity of FIE mRNA after 96h of 10 uM estradiol treatment vs mock (DMSO), normalized to 0h mock treated samples, 3 biological replicates. (**D**) Relative quantity of PHT1:2 and PHT1:9 mRNA before (0h) during (4h and 96h) and after (1h and 24h) of -P starvation. Plants were germinated and grown for 10 days on ½ MS media with 10 uM estradiol or “mock’ plates (same media but with DMSO).

Based on our understanding of gbH2A.Z and H3K27me3 as barriers to transcriptional activation, it was surprising to observe both relatively unchanged despite a vast transcriptional reprogramming during phosphate starvation and resupply. While H3K27me3 does not appear to be dynamic in response to phosphate starvation or repletion, many PSi genes are highly enriched for H3K27me3 (**Figure 4b**). To assess the dependence of the PSi response on H3K27me3, we exposed PRC2 mutant plants (*clf-28* homozygotes) to a P starvation and resupply event (**Figure S4**). Using RT-qPCR, we found no significant change in induction or silencing rates of three canonical polycomb-target PSi genes in *clf-28* mutants compared to WT (**Figure S4**). However, it is possible that the loss of CLF is compensated by expression of a paralogous SET domain PRC2 component, SWINGER. To account for this possibility, we developed an estradiol inducible artificial microRNA (amiRNA) knockdown of *FERTILIZATION INDEPENDENT ENDOSPERM* (*FIE*), a nonredundant and essential component of the PRC2 complex (Chanvivattana et al. 2004). After confirming efficient knockdown of FIE after 10 days of amiRNA induction (**Figure 5c**), we put plants through a phosphate starvation and resupply event and measured expression of the *PHT1;2* and *PHT1;9* PSi genes by RT-qPCR in roots. Surprisingly, we observed that PSi gene induction and repression after resupply are not affected by this loss of PRC2 activity, unlike nitrogen and drought stress response genes (**Figure 5d**) (Bellegarde et al. 2018; Liu et al. 2014). There is a slight decrease in expression of *PHT1;9* and *PHT1;2* when comparing the *FIE* knockdown plants to controls. However, these differences are slight and the rate of induction, absolute expression, and repression are not significantly affected.

### Genes induced after 30m of P resupply lose H2A.Z and H3K4me3 while genes induced after 96h of P starvation largely do not

We initially included H3K4me3 to use in combination with RNA-seq to distinguish post-transcriptional changes in RNA abundance from increases in transcription. Interestingly, we found that the relationship between RNA levels and H3K4me3 levels was positively correlated over long time scales but negatively correlated on shorter time scales (**Figure 3c**). After 96h of -P stress, the expression of genes gaining H3K4me3 increased significantly more than those without H3K4me3 changes or those losing H3K4me3 (**Figure 6a**). These results follow the expected trend and suggest that increased H3K4me3 is not absolutely necessary for activation, but may facilitate the degree of gene activation in some cases.

**Figure 6.**
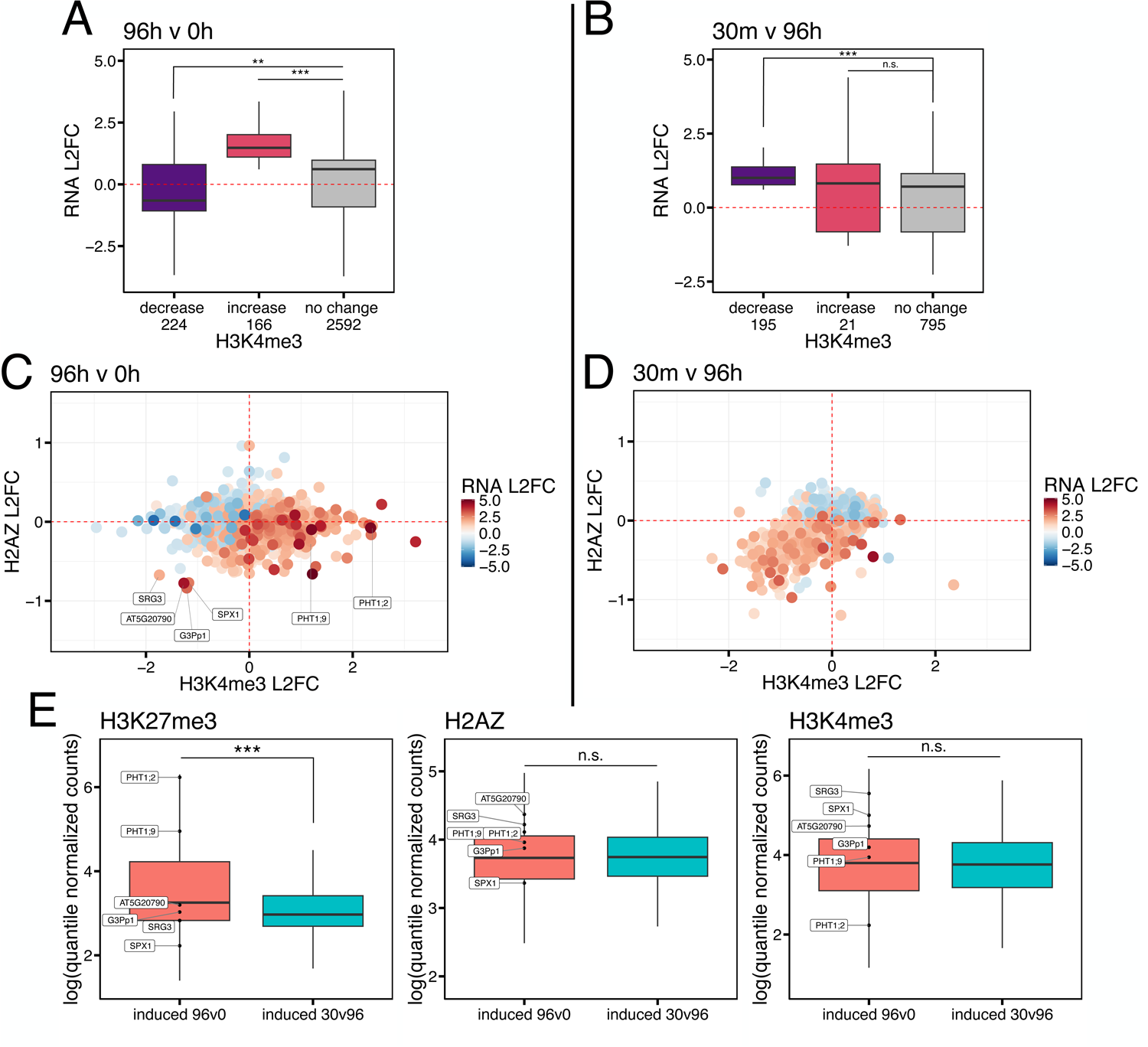
Genes induced after 30m of P resupply lose H2A.Z and H3K4me3 while genes induced after 96h of P starvation do not. (**A**) Boxplot of RNA L2FC of genes with increased, decreased and unchanged H3K4me3 after 96h of Pi starvation. ***p<0.001 (unpaired Student’s t-test) (**B**) Same as (A) for RNA L2FC from 96h Pi starvation to 30m Pi resupply. (**C**) L2FC of H3K4me3 after 96h of P starvation vs L2FC of H2A.Z after 96h of P starvation. Color represents L2FC of RNA after 96h of starvation. Points represent all differentially expressed genes from 0h to 96h -P (|L2FC| > 1, padj < 0.05). (**D**) Same as C for L2FC from 96h P starvation to 30m Pi resupply. (**E**) Quantile normalized counts at genes induced after 96h -P vs genes induced after 30m +P. ***p<0.001 (Wilcoxon two-sided test).

We next extended this analysis to time points after P resupply and noticed that the relationship between H3K4me3 and RNA dynamics is largely reversed in the 30m after P resupply (**Figure 6b**). At 96h of P starvation, only 48% (114) of differentially expressed (DE) genes that lose H3K4me3 are induced while 93% (196) of DE genes that gain H3K4me3 are induced. This is a dramatic difference compared to H3K4me3 dynamics 30m after P resupply, where 91% (181) of DE genes that lose H3K4me3 are induced while only 64% (16) of DE genes that gain H3K4me3 are induced. Interestingly, of the 29 genes that become induced and lose H2A.Z after 30m +P, 23 (79%) also lose H3K4me3 at the same time. This appears to occur without any changes to chromatin accessibility or H3K27me3, with both features barely changing from 96h -P to 30m +P (**Figure S6**).

We finally expanded our analysis to all differentially expressed genes irrespective of their chromatin changes. We compared the L2FC of H2A.Z to the L2FC of H3K4me3 at all differentially expressed genes after 96h of P starvation and again after 30m of P resupply. After 96h of P starvation, gene activation is poorly correlated with H2A.Z changes and has a weak positive correlation with H3K4me3 changes (**Figure 6c)**. Interestingly, this relationship is flipped after 30m of +P resupply (**Figure 6d**). Here, H3K4me3 and H2A.Z are anticorrelated with increases in transcript, and the correlation is much stronger than after 96h -P, with most induced genes losing H2A.Z and H3K4me3 (R^2^ = 1.69E-7 at 96h v 0h, R^2^ = 0.277 at 30m v 96h). One possible explanation for these results is that the early stages of transcriptional induction (e.g. 30 min + P) are more disruptive to nucleosomes, causing losses of H2A.Z/H2B and H3/H4 dimers, which may be ameliorated after longer periods of induction (e.g. 96 h - P), allowing stabilization of H2A.Z levels and accumulation of H3K4me3.

We noticed that a small group of genes induced after 96h -P stand out by significantly losing H2A.Z and H3K4me3, similar to the majority of induced genes after 30m P resupply (**Figure 6c**, SPX1, SRG3, G3pP1, and AT5G20790). Intriguingly, these same genes were previously found to be the four highest expressed PSi genes after 30m of P starvation in whole root tissue [20]. Perhaps the mechanism of activation observed after 30m of Pi resupply is sustained in these 4 genes even after 96h of starvation. Interestingly, we found that at steady state, there is significantly less H3K27me3 at genes induced after 30m of P resupply than genes induced after 96h of P starvation, while H2A.Z and H3K4me3 levels were comparable (**Figure 6e**). Additionally, the 4 genes that do lose H2A.Z and H3K4me3 after 96h -P had H3K27me3 enrichment levels similar to the genes induced after 30m (**Figure 6e**). This raises the notion that H3K27me3 may aid in retention of nucleosomal components during transcription.

## Discussion

In this study we used INTACT cell type-specific nuclei purification followed by RNA-seq, ATAC-seq, and CUT&Tag to profile the transcriptomic and epigenomic responses of the Arabidopsis root hair cell type to a time course of phosphate starvation and subsequent resupply (**Figure 1a**). Analysis of our nuclear RNA-seq data across the time course identified 6 major patterns of transcript level changes. Time course cluster 3 (TC3) contained transcripts increased by P starvation that returned to baseline levels after 4 hours of P resupply (**Figure 1c**). The TC3 class represents classical phosphate starvation-induced (PSi) transcripts, whose regulatory regions were found to be enriched for binding sites of TFs known to drive the PSR (**Dataset S3**). We also identified two classes of phosphate starvation-repressed (PSr) transcripts in the TC5 and TC2 groups. The behavior of these classes of PSr transcripts differ in that TC5 transcripts return to pre-starvation levels after 4 hours of phosphate repletion, while those in the TC2 class do not. Interestingly, there are 3 classes of transcripts that respond distinctly to resupply of P after starvation. TC1 class transcripts reach a maximum level after 30 minutes of resupply and return to baseline within 4 hours. The TC4 genes show a very similar profile but are only induced after 30 minutes of resupply and reach a maximum level at 1 hour before returning to baseline at 4 hours. Finally, the TC6 class transcripts do not show significant alterations during P starvation and generally decrease progressively in the 4 hours after resupply. In total, we identified 5,674 differentially expressed genes between any two time points across the time course, indicating major reprogramming in the root hair cell transcriptome. While some of the transcript level changes are likely due to post-transcriptional control mechanisms, others are driven by changes in transcription. These data offer an opportunity to explore a variety of questions and could be used in conjunction with other datasets, for example a previous root hair nuclear RNA secondary structure and RNA-binding profile analysis (Foley et al. 2017).

To address the relationships between transcript levels and chromatin features, we employed ATAC-seq as well as CUT&Tag to monitor histone variant H2A.Z, the transcription-associated mark H3K4me3, and polycomb silencing-associated mark H3K27me3. Comparison of our root hair CUT&Tag datasets for each mark with those generated using ChIP-seq on seedling tissue showed a strong concordance in locations and profiles of each signal (**Figure 2a-c**), giving confidence that our data are accurate. Major advantages of the CUT&Tag approach compared to ChIP-seq are the low nuclei input requirement and higher-signal to noise ratio, which allowed us to manageably generate triplicate datasets for each mark across 6 time points. Combining root hair RNA-seq data with chromatin profiles under phosphate-replete conditions, we found that transcript levels correlate well with each chromatin profile at steady state (**Figure 2d**). To our surprise, however, the chromatin changes associated with differential transcript levels during phosphate starvation and recovery were relatively minimal across the time course and were seen most prominently for H3K4me3 (**Figure 3b**). Of the chromatin changes that were significant after 96 h -P, many were retained after 4 hours of phosphate resupply, when the corresponding transcripts had returned to baseline levels (**Figure 3a**). For example, 80% of the genes significantly altered in H3K4me3 levels after 96 h -P were still differentially enriched at 4 hours after resupply. The relatively limited chromatin changes observed relative to transcript level changes is likely due in part to post-transcriptional regulation mechanisms.

A major motivation for this study was to understand how polycomb-target genes marked with both H3K27me3 and gene body H2A.Z are activated and re-silenced. We thus focused on genes induced by 96 h -P (the TC3 class), as many of these PSi genes were already known targets of PRC2 and H2A.Z. Examination of the starting chromatin state of these and each of the other 5 DEG clusters confirmed that the TC3 genes were highly enriched for H3K27me3 and gbH2A.Z, although not exclusively. Roughly 30% of these TC3 genes showed very low or no expression, low chromatin accessibility, gbH2A.Z and high H3K27me3 levels (**Figure 4a** and **b**, **Figure 5a** and **b**). While these genes showed large gains in transcript levels during P starvation, they surprisingly showed minimal changes to H3K27me3 and H2A.Z (**Figure 4c**). This is counter to findings in similar experiments exploring H2A.Z dynamics which showed pronounced H2A.Z loss at genes induced during drought, heat, osmotic, hypoxic, light stress, and ethylene responses (Boden et al. 2013; Mao et al. 2021; Cortijo et al. 2017; Nguyen and Cheong 2018; Sura et al. 2017; Lee and Bailey-Serres 2019; Zander et al. 2019).

The lack of a requirement for H2A.Z eviction during transcriptional induction was also recently observed in our lab when profiling the dynamics of H2A.Z and the SWR1 complex at gbH2A.Z/H3K27me3-enriched ABA responsive genes (Krall and Deal 2024). While many of these genes were induced following 4h of ABA treatment, H2A.Z levels and distribution across most were unperturbed or only slightly diminished. Interestingly, induction of these previously silent genes by ABA was accompanied by proportional increases in recruitment of SWR1 complex components, suggesting that SWR1 is recruited to replace H2A.Z lost during transcription. Thus, it seems likely that a similar mechanism is operating at PSi genes to maintain gbH2A.Z during transcription. The question remains open as to how these genes become permissive for transcription while retaining gbH2A.Z and H3K27me3..

Some H3K27me3/gbH2A.Z genes, such as those responsive to temperature, far-red light, and ethylene appear to depend upon H2A.Z removal by the INO80 complex for induction (Willige et al. 2021; Zander et al. 2019; Zhao et al. 2023; Xue et al. 2021), and this may be coupled with active H3K27me3 removal by REF6 (Zander et al 2019) or perhaps passive loss due to H2A.Z eviction. In contrast, the PSi genes described here lose neither mark yet are able to support transcriptional induction. How this happens is not yet clear but could involve simple loss of ubiquitinated H2A.Z (Gómez-Zambrano et al. 2019) from chromatin in early rounds of transcription and replacement with unmodified H2A.Z by SWR1, and/or active deubiquitination by enzymes such as UBP5 (Godwin et al. 2024). The emerging picture is one of multiple mechanisms for activation of polycomb/gbH2A.Z target genes, where in some cases H2A.Z is evicted, and others where it is actively replaced.

Given the unusually high levels of H3K27me3 at polycomb/H2A.Z target genes in TC3 (**Figure 5a** and **b**), we speculated that while H3K27me3 may not be dynamic across the activation/repression of these genes in response to phosphate, H3K27me3 could affect the kinetics of induction or repression, or the absolute levels of transcript produced upon induction. *CURLY LEAF* deficient plants were previously shown to have an aberrant increase in expression of *NRT1.2* during nitrogen starvation (Bellegarde et al. 2018). A similar phenomenon was observed in drought stress response genes, where H3K27me3 remained unchanged during drought stress at induced genes but *clf* mutants displayed higher induction of drought responsive genes under stress (Liu et al. 2014). Counter to these findings, we saw no such ablation of the rate of induction, absolute expression, or repression of H3K27me3 enriched PSi genes *PHT1;2*, *PHT1;9,* or *ASK11* after P stress and resupply in *clf* or an estradiol inducible knockdown of the PRC2 core subunit *FIE* (**Figure 5c** and **d, Figure S4**). Interestingly, a recent study of cold stress in Arabidopsis also observed limited H3K27me3 changes upon gene induction, and a lack of requirement for CLF in maintaining gene repression (Faivre et al. 2024). Taken together, the differences observed in gene regulation between environmental responses argue against a single model of H2A.Z and H3K27me3 action and invite further investigation through direct comparisons of different responses.

Another interesting and unexpected finding in this study was the shift in correlations between transcript levels and those of H2A.Z and H3K4me3 across different time points. Considering all differentially expressed genes after 96h of -P, H3K4me3 changes show a positive correlation with transcript level changes, but this relationship is reversed when comparing the 96h -P time point to the 30 min resupply time point (**Figure 3c**). This trend becomes clearer upon examination specifically of genes with differential H3K4me3 enrichment across these two different time windows (**Figure 6a** and **b**). Interestingly, we observed that DEGs after 96h -P tend to show an increase in enrichment of H3K4me3 when induced and a reduction of H3K4me3 when repressed, with very little change in H2A.Z regardless of the direction of change (**Figure 6c**). In contrast, genes induced in expression after 30 minutes of P resupply show a loss of both H3K4me3 and H2A.Z (**Figure 6d**). Both changes after 30 m proved to be transient, as of the 22 genes that significantly lost H2A.Z and 2764 genes that significantly lost H3K4me3 after 30m of P resupply, only 1 and 313 were still differentially enriched by 4h after P resupply, respectively. A distinguishing feature of the genes induced after 96 h -P is a higher level of H3K27me3 compared to those induced after 30 min of P resupply. The differences in chromatin dynamics observed between the gene sets could thus reflect a role for H3K27me3 in promoting the retention of nucleosome components during transcription. Given the transient loss of H2A.Z at induced genes after 30m of resupply, and the new finding that SWR1 enrichment increases at induced genes (Krall and Deal 2024), we propose that loss of H2A.Z at 30m +P induced genes is quickly replenished by SWR1. Therefore, a sample of the transcriptome at a later time (4h+P) would only capture increased transcript levels but a chromatin sample taken at the same time would not capture the corresponding H2A.Z loss. This phenomenon could explain the 4 outlier PSi genes that lose H2A.Z after 96h (**Figure 6C**). In this scenario, H2A.Z incorporation by SWR1 would not occur fast enough to account for H2A.Z lost via transcription, thus a loss of H2A.Z is observed. These results indicate that distinct relationships between transcription and chromatin dynamics can exist, and these may be time-dependent with respect to the onset and level of transcription at a given gene.

In conclusion, here we report several surprising findings with respect to the relationships between transcription and chromatin changes on short time scales and in the absence of cell division, which we hope will spur further work. In addition, the data generated here will serve as a useful resource for the field to explore how P starvation response and recovery are orchestrated in the root hair cell type, as well as to probe additional questions about the complex relationships between transcriptional activity and the epigenome.

## Methods

### Growth conditions and transformation

Seeds of Arabidopsis thaliana (Col-0 strain) were grown on half-strength Murashige and Skoog (½ MS) media agar plates supplemented with 1% sucrose. Plates were stratified in dark conditions for 3 days at 4°C followed by 10 days of vertical growth at 20°C under a 16 hour light/8 hour dark cycle. Plasmids used for plant transformation were introduced into *Agrobacterium tumefaciens* GV3101 strain by electroporation and transformed into plants using the floral dip method (Clough and Bent 1998). Transgenic plants were selected on ½ MS containing 50 mg/L Kanamycin and 100 mg/L timentin.

For phosphate starvation conditions, seedlings were transferred to ½ MS media without potassium phosphate and supplemented with 1% sucrose and potassium sulfate to balance the ionic composition with phosphate rich plates. After 96 hours of starvation, seedlings were transferred back to ½ MS containing phosphate.

### Plasmid DNA constructs

To produce an INTACT construct driven by the root hair-specific *ADF8* promoter (AT4g00680) we first cloned the *AtADF8* regulatory region from - 750 bp to +427 bp relative to the TSS into the gateway-compatible pENTR-D/TOPO plasmid (Invitrogen). This fragment consisted of the promoter, first exon (one codon), first intron of ADF8, which contains a root hair-specific enhancer, as well as the first two codons of exon 2. We then subcloned this fragment into pK7WG-INTACT-At gateway destination plasmid (Ron et al. 2014) using the LR clonase II enzyme in an LR recombination reaction (Invitrogen) so that the NTF component of INTACT was driven by the *ADF8* regulatory sequences. The amiRNA construct targeting the first exon of *FIE* (GTCTTCGTTACCGCTGGTGGA) was generated with zero mismatches using the “Design” feature of the Web MicroRNA Designer (WMD3; Schwab et al. 2006) and synthesized into a Gateway entry vector by TWIST Bioscience. The construct was then recombined by Gateway cloning into pMDC7 for estradiol inducible expression and pEG100 for constitutive expression via a 35S promoter (Earley et al. 2006; Curtis and Grossniklaus 2003).

### Real-time RT-PCR (qRT-PCR) for testing PSi genes in *clf-28* and FIE knockdowns

Total RNA was isolated from root tissue 1 cm above the root tip using the RNeasy plant mini kit (Qiagen). Extracted RNA was DNase treated and converted into cDNA with LunaScript RT SuperMix kit (New England Biolabs). cDNA was used as templates for real-time PCR on StepOnePlus real-time PCR system (Applied Biosystems) using SYBR Green as a detection reagent. *PP2A* mRNA (AT1G13320) primers were used as the endogenous control (Czechowski et al. 2005). The primer sets used can be found in **Supplemental Table 1**.

### RNA-seq library preparation and analysis

Biotin tagged nuclei from ADF8 INTACT root tissue (∼100 mg) were isolated from a 1 cm section of the fully differentiated root hair zone (above the differentiation zone and below the first lateral root) and captured on 25 uL of Streptavidin M280 Dynabeads according to the Isolation of Nuclei TAgged in specific Cell Types (INTACT) protocol (Bajic et al. 2018). Bead bound nuclei from each timepoint in the starvation assay were counted and nuclei were split three ways to ∼100,000 nuclei per sample in NPBt for three replicates. RNA was isolated from bead bound nuclei using the RNeasy plant micro kit (Qiagen) with on column DNase treatment and quantified via Quant-it™ RiboGreen RNA Assay Kit (Thermo-Fisher). Libraries for RNA-seq were generated according to the QuantSeq 3’ mRNA-Seq Library Prep Kit FWD V1 (Lexogen, cat.no. 015.24) following the manufacturer’s protocol with modified steps for low input samples.

All amplified libraries were pooled and sequenced using single-end 50 nt reads on an Illumina NextSeq 2000 instrument. Reads were mapped to the *Arabidopsis thaliana* TAIR10 genome assembly, converted to the BAM file format, and sorted by coordinate using the STAR aligner with the default parameters (Dobin et al. 2013). Duplicate reads were removed using Picard markDuplicates (Broad Institute 2019). Strand specific fragment counting on genes was performed with a TAIR10 specific GTF file using featureCounts with the following parameters: -s 1 -t exon -g gene_id (Liao et al. 2013). Differentially expressed genes were clustered based on direction of response to phosphate starvation using TCseq (Mengjun 2019). Briefly, pairwise differential analysis was performed across all time point combinations using DBanalysis. Then results were filtered for genes with an adjusted p value >0.05 and absolute fold change > 2 using timecourseTable. Finally, the remaining differentially expressed genes were grouped into 6 c-means clusters using timeclust. Gene ontology was performed via WEBGestalt using default parameters (Elizarraras et al. 2024). Simple Enrichment Analysis was performed on all TC3 gene promoters overlapping with ATAC hypersensitive sites using the SEA tool in the MEME Suite with default parameters and all other accessible TC gene promoters as a control (Bailey and Grant 2021). To find a control set of equally expressing genes, average RNA counts at 0h for each TC3 gene were matched to a gene with equivalent average RNA counts at 0h that was not differentially expressed.

### INTACT-CUT&Tag and ATAC-seq on root hair nuclei

Biotin tagged nuclei from ∼100 mg from ADF8 INTACT root tissue (same 1 cm segment as for RNA-seq) were captured on 25 uL of Streptavidin M280 Dynabeads according to the Isolation of Nuclei TAgged in specific Cell Types (INTACT) protocol (Bajic et al. 2018). Bead bound nuclei for each timepoint in the starvation assay were counted and nuclei were split three ways to ∼100,000 nuclei per sample in NPBt for three replicates.

CUT&Tag was performed on these bead bound nuclei (100,000 per reaction) according to the Epicypher CUT&Tag Protocol v1.7 (EpiCypher 2022) with the following changes. All buffers were prepared without Digitonin and streptavidin bead bound nuclei were resuspended directly from NPBt into 50 uL of Antibody150 Buffer containing 0.5 ug of anti-H3K27me3 (Millipore 07-449), anti-H2A.Z (Deal et al. 2007), anti-H3K4me3 (Milipore 07-473) or anti-IgG (sigma I-5006) and incubated at 4°C overnight. Guinea pig anti-rabbit secondary (0.5 ug, Sigma SAB3700889) was incubated for 1 hr at room temperature in 50 uL Wash150 Buffer. Tagmented libraries were amplified using 17 PCR cycles. ATAC was performed in duplicate (∼20k nuclei each) on the same pools of bead bound nuclei, as previously described (Bajic et al. 2018).

### INTACT-CUT&Tag sequencing and data analysis

All amplified libraries were pooled and sequenced using paired-end 150 nt reads on an Illumina NovaSeq 6000 instrument. Data processing was performed with guidance from Zheng et al. 2020 (Zheng et al. 2020). Reads were trimmed of adapter content using Trimgalore and mapped to the *Arabidopsis thaliana* Col-PEK genome assembly (Hou et al. 2022) using Bowtie2 with the following parameters: --local --very-sensitive --no-mixed --no-discordant --phred33 -I 10 -X 700 (Krueger). Aligned reads were converted to the BAM file format using Samtools (Li et al. 2009). Duplicate reads were removed using Picard markDuplicates (Broad Institute 2019). For CUT&Tag samples, reads from fragments less than 150bp were discarded to eliminate sub-nucleosome sized fragments. Peaks were called on deduplicated BAM files using MACS2 with IgG as a control where applicable (Feng et al. 2012). For visualization, deduplicated BAM files of each sample were RPKM normalized and subtracted from their corresponding normalized IgG BAM file using bamCompare from deepTools (Ramírez et al. 2014). RNA, ATAC, and CUT&Tag average plots stratified by expression quantile were generated using SeqPlots (Stempor and Ahringer 2016). Quantile normalized counts for each CUT&Tag result were generated using multibigwigsummary with --binSize 100 to generate a matrix of counts across samples (Ramírez et al. 2014). The resulting matrix was quantile normalized using the R package preprocessCore. Quantile normalized scores per 100 bp bins were then averaged across single genes and again across biological replicates. Mapped fragments overlapping with the TSS (−200bp +100bp from TSS) were counted using featureCounts for ATAC samples (Liao et al. 2013). Mapped fragments overlapping with genes were counted using featureCounts for CUT&Tag samples. Gene counts were then analyzed for differential enrichment using DESeq2 (Love et al. 2014). DESeq2 results were visualized in R using ggplot2 and pheatmap packages (R Core Team 2024).

### RT-PCR and ChIP-qPCR to confirm RNA-seq and Cut&Tag results at *PHT1;2* and *PHT1;9*

Wild type seeds were germinated on ½ MS media and grown vertically for 5 days before seedlings were transferred to ½ MS without Pi. Whole root samples were taken just before transfer to -P media and 48 hours after transfer to -P media. RNA was isolated using the Qiagen RNeasy Plant Mini Kit and first strand cDNA was produced using the SuperScript III first-strand cDNA kit (Thermo). cDNAs from the +P and -P samples were used as templates for real-time PCR on the StepOnePlus real-time PCR system (Applied Biosystems) using SYBR Green as a detection reagent and the ACTIN2 transcript as an endogenous control.

Samples for ChIP-qPCR were crosslinked with formaldehyde before processing using the enhanced ChIP protocol reported previously (Zhao et al. 2020), with the following modifications: per replicate, 200 mg of 7-day old root tissue was harvested, crosslinked in 1% formaldehyde, and flash frozen in liquid nitrogen. Tissue was ground in a mortar and pestle, homogenized into a slurry in 250 µl of Buffer S and moved directly to sonication. Post sonication, lysate was diluted 10x with Buffer F for use in pulldowns. 50 µl of diluted lysate was saved as input. Antibodies against H2A.Z (2 μg/mL) (described above for CUT&Tag) or αH3K27me3 (0.5 μg/mL) (Millipore 07-449) antibody was added to diluted lysates and incubated overnight at 4°C on a nutator. 20 µl of washed Protein G Dynabeads (Invitrogen) was added to each pulldown. Antibodies against H2A.Z and H3K27me3 were the same as used for CUT&Tag. Two biological replicates were performed using the 3’ UTR of the ACT2 gene as the endogenous control for both RT-qPCR and ChIP-qPCR. The primer sets used can be found in **Supplemental Table 1**.

### Publically available data

Seedling ChIP-seq data for H3K27me3, H2A.Z, and H3K4me3 (Figure 2a) were obtained from the NIH Sequencing Read Archive database under the following accessions, respectively; SRR5278091, SRR5364421 and SRR5364426 (Zhou et al. 2017; Wollmann et al. 2017). Data was processed as above.

## Data Availability

All sequencing data generated in this study have been deposited in the NCBI GEO database under accession number GSE271993.

---

**Supplemental Figure 1.**
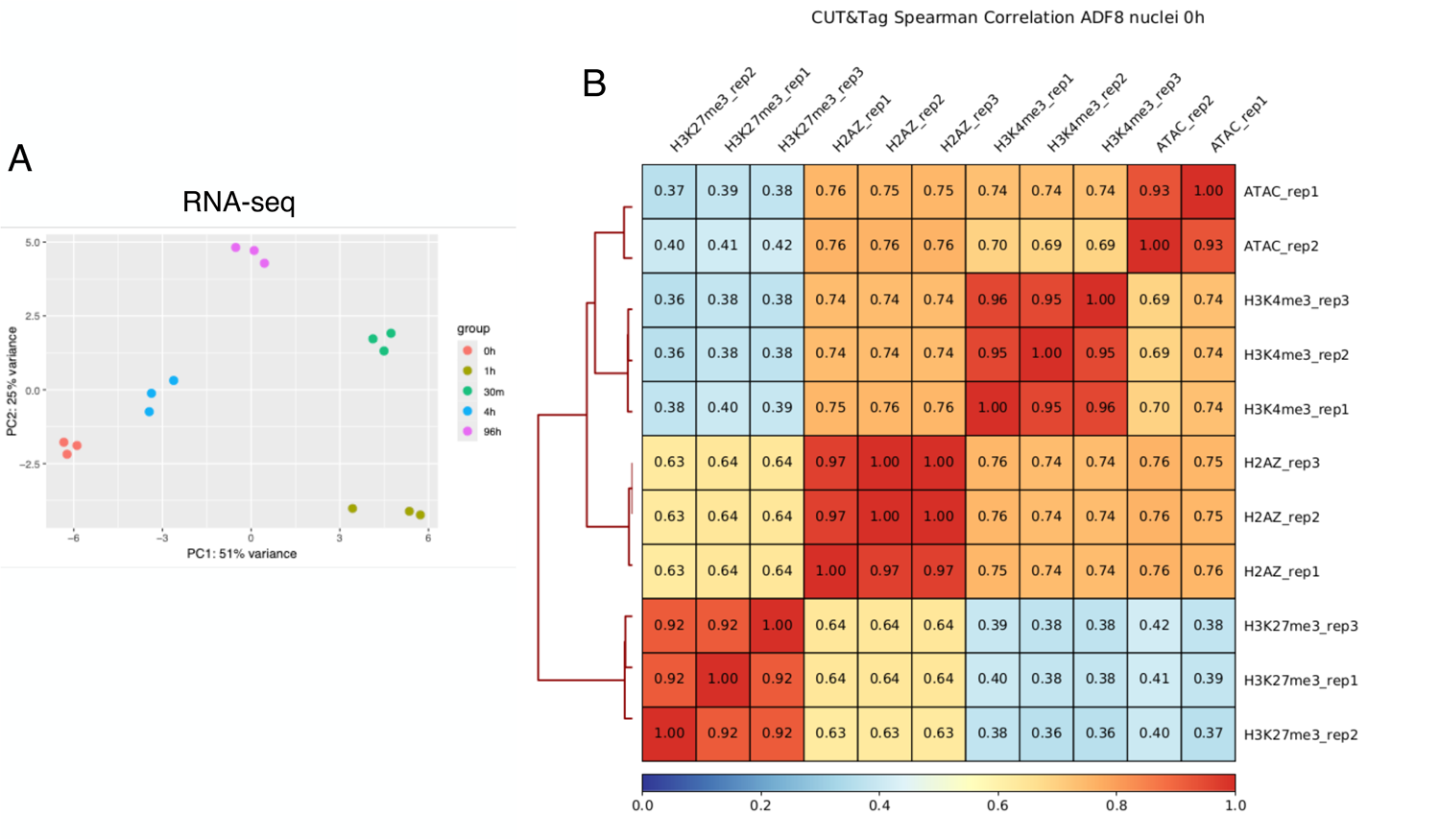
Overview of RNA-seq and chromatin profiling data. (A) PCA1 vs PCA2 of RNA DESeq2 results across all timepoints (3 biological replicates per time point). (**B**) 1000 bp bin genome wide hierarchical clustered Spearman correlation across every CUT&Tag and ATAC-seq replicate at 0h.

**Supplemental Figure 2.**
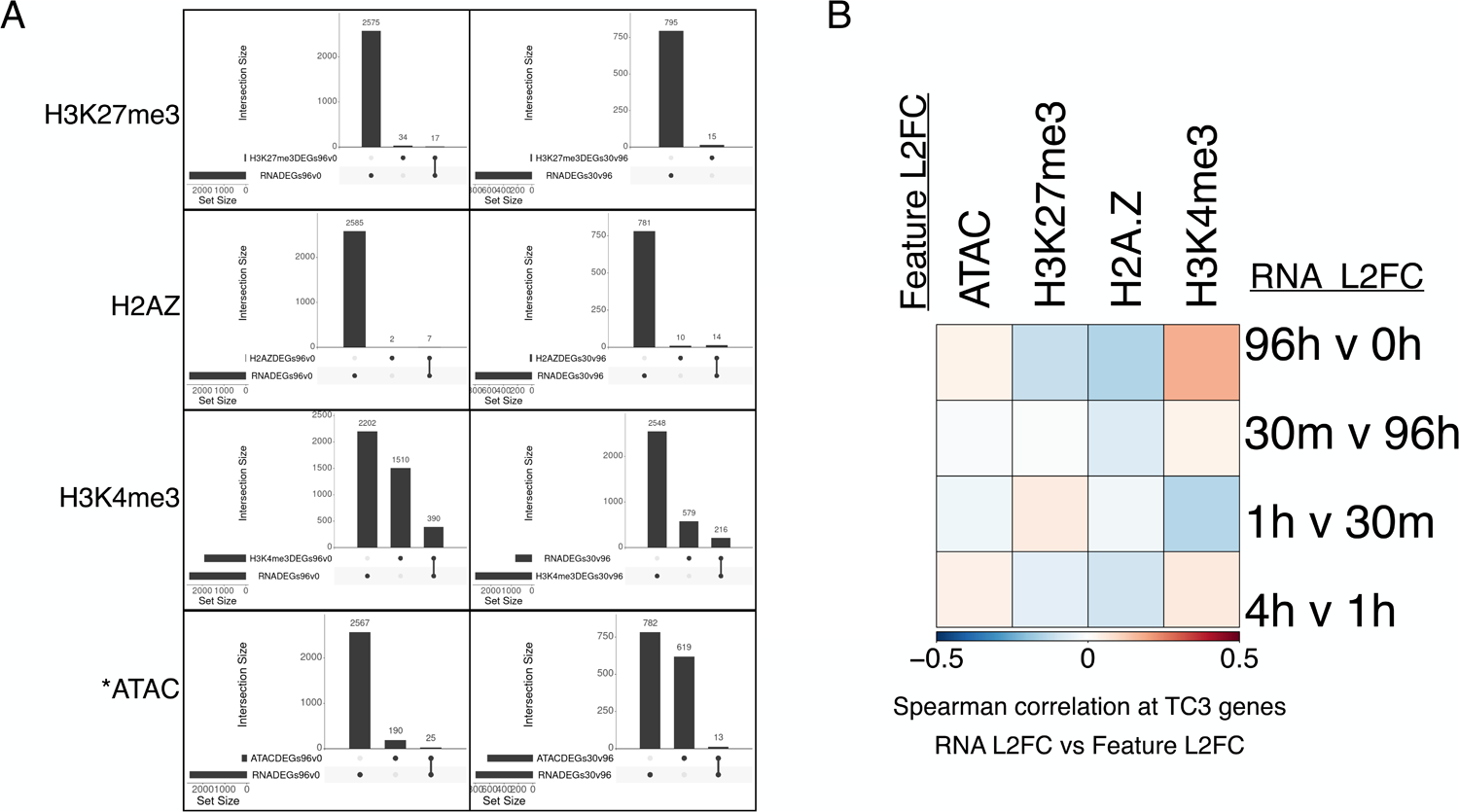
Chromatin features across the time course. **(A)** Upset plot of overlap between differentially enriched chromatin features (H3K4me3, H3K27me3, H2A.Z) and differentially expressed genes at 96h vs 0h and 30m vs 96h. *Differentially accessible promoters defined by ATAC were determined via p-value < 0.05 due to only one gene being significantly different when determined by the typical adjusted p-value. (**B**) L2FC Spearman correlation between RNA and each chromatin profile across the time course exclusively at TC3 genes. Each L2FC is measured relative to the previous time-point.

**Supplemental Figure 3.**
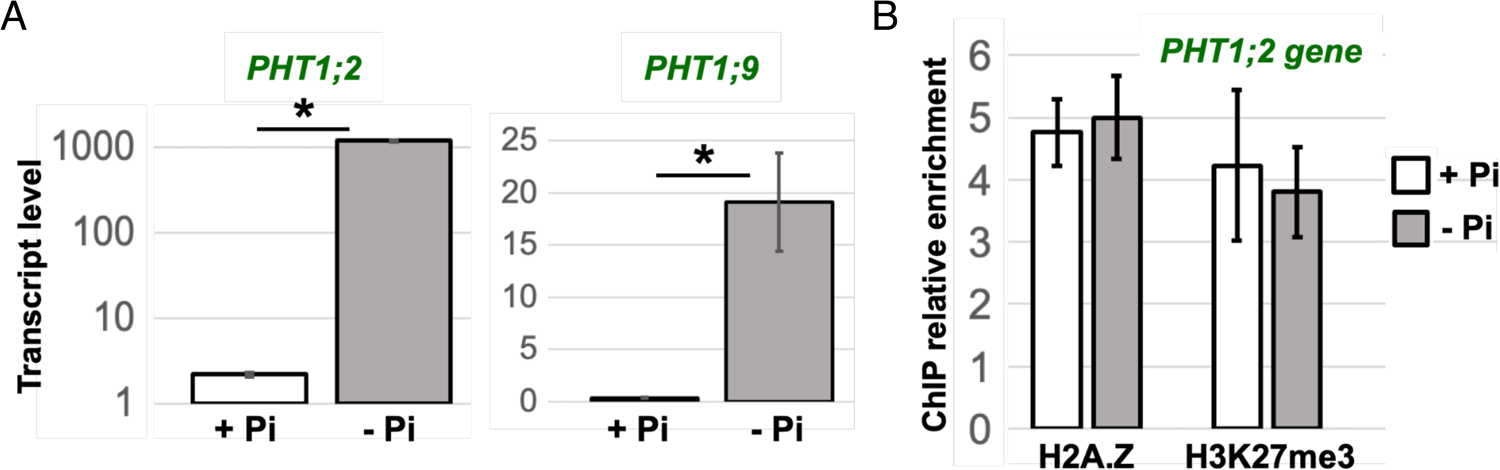
Confirmation of transcript, H3K27me3, and H2A.Z levels at PSi genes in whole root. (**A**) Quantitative RT-PCR analysis of RNA from whole roots grown on +P media and transferred to -P media for 48 hours. Error bars represent standard deviation from the mean for 3 replicates. (**B**) ChIP-qPCR analysis of H2A.Z, H3K27me3, and H2AUb enrichment at *PHT1;2* relative to ACT2 control site under + and -P conditions. Error bars represent standard deviation from the mean for 3 replicates.

**Supplemental Figure 4.**
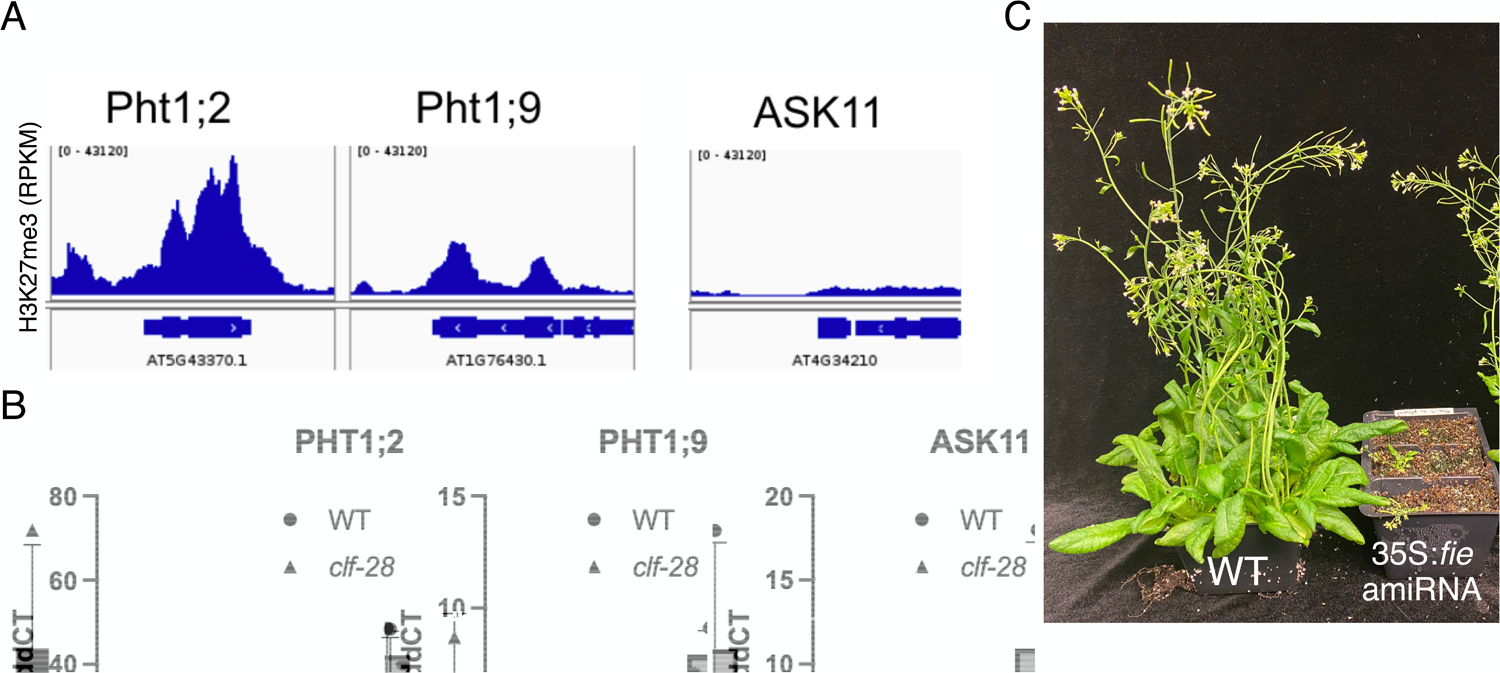
Phenotypes of PRC2 knockdown plants. (**A**) RPKM normalized H3K27me3 enrichment at three PSi genes. (**B**) Quantitative RT-PCR analysis of RNA from whole roots subject to P starvation and repletion gray bars represent clf-28 mutant plants. Error bars represent standard deviation from the mean for 3 replicates. Students t-test found no significant difference between WT and clf-28 at any gene. (**C**) WT vs 35S:fie amiRNA knockdown of PRC2 subunit FIE after 1 month growth on soil.

**Supplemental Figure 5.**
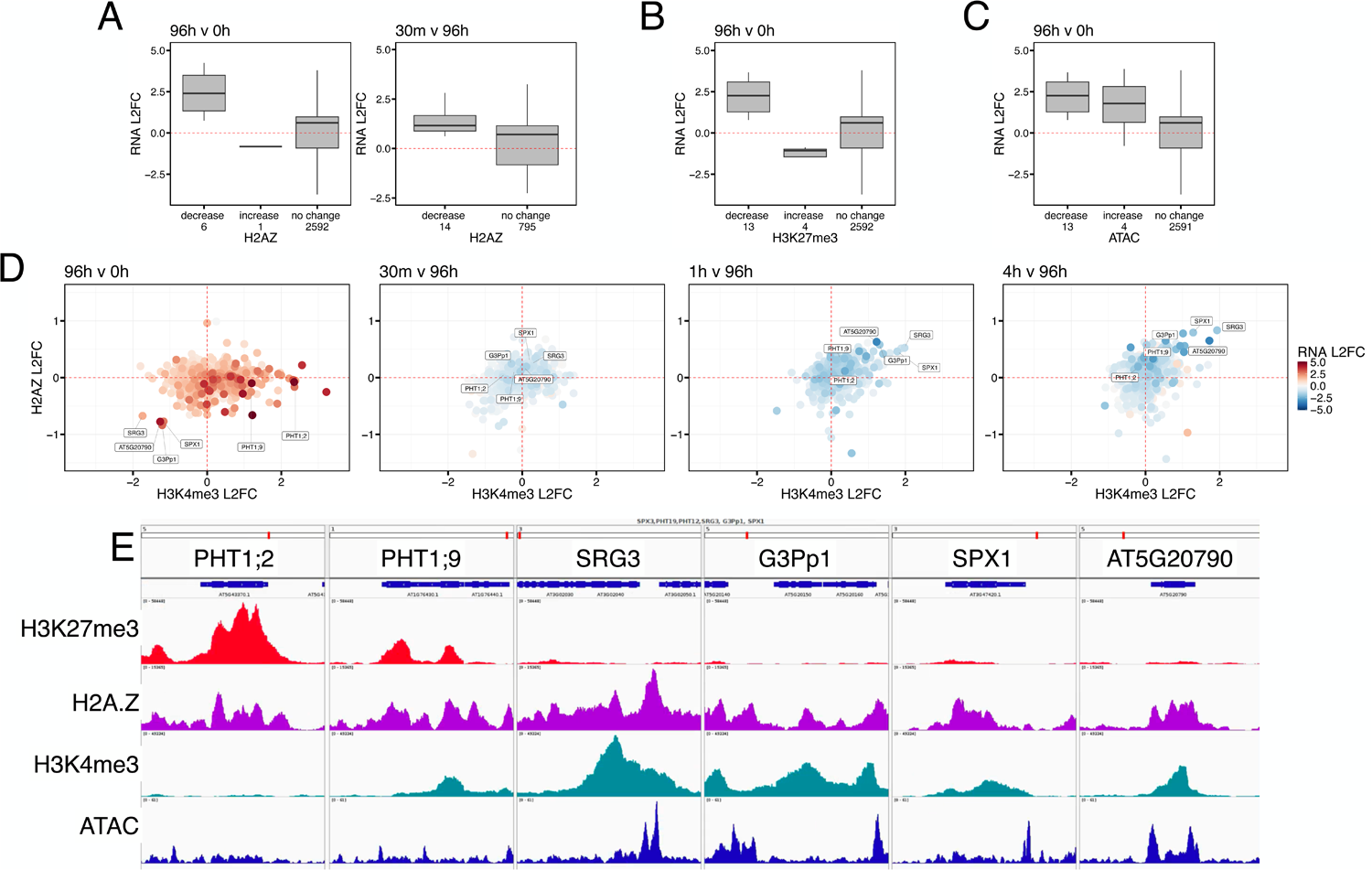
Expression changes at genes with differential chromatin enrichment, chromatin dynamics of PSi genes across the starvation time course and 0h chromatin profiles at genes of interest. (**A,B,C**) Boxplot of RNA L2FC of genes with increased, decreased and unchanged H2A.Z, H3K27me3, and ATAC after 96h of Pi starvation and after 30m of resupply (no DE genes for H3K27me3 or ATAC after 30m). (**D**) L2FC of H3K4me3 after 96h of P starvation vs L2FC of H2A.Z relative to the most recent P change. Color represents L2FC of RNA. Points represent all TC3 genes. (**E**) RPKM normalized enrichment for PSi genes of interest from TC3.

